# Low Shear Modeled Microgravity Induces Unexpected Motility Phenotypes in *Salmonella* Typhimurium

**DOI:** 10.64898/2026.06.18.731987

**Authors:** Jiseon Yang, Jennifer Barrila, Laura Banken, Karla P. Franco Meléndez, Christian L. Castro, Bianca Y. Kang, Sandhya Gangaraju, Richard R. Davis, C. Mark Ott, Robert JC McLean, Cheryl A. Nickerson

## Abstract

Bacteria routinely exhibit unexpected phenotypic and molecular changes in response to spaceflight and spaceflight-analogue conditions, yet the mechanisms by which they sense and respond to these low fluid shear environments are not fully elucidated. We previously demonstrated that spaceflight and low shear modeled microgravity (LSMMG) altered motility and chemotaxis gene expression in *Salmonella enterica* serovar Typhimurium (*S*. Typhimurium), raising the possibility that flagella mediate responses of the pathogen to these environments. Herein, we investigated whether LSMMG culture alters *S*. Typhimurium motility and examined the role of flagella in regulating pathogenesis-associated stress and infection phenotypes. LSMMG enhanced the swimming motility of wild-type *S*. Typhimurium relative to 1x*g* controls; a trend which persisted even in the absence of the global stress response regulators Hfq and RpoS. This finding was unexpected, as Δ*hfq* mutants are typically defective for motility under conventional culture conditions. Motility was also observed in the flagella-deficient Δ*flhDC* mutant following LSMMG and 1x*g* culture, although the relative motility pattern differed relative to wild-type. Collectively, these results indicate that flagella contribute to LSMMG-enhanced motility, but are not strictly required under these conditions. Conditioned supernatant exchange demonstrated that LSMMG-induced motility changes are cell-intrinsic rather than mediated by extracellular factors. While flagella were dispensable for many pathogenesis-related phenotypes tested, their deletion selectively altered the magnitude of LSMMG-associated thermal stress and intracellular survival in human intestinal epithelial cells. Together, these findings demonstrate that motility and pathogenesis-related responses in *S. Typhimurium* are governed by multiple regulatory pathways that differentially respond to LSMMG and 1x*g* conditions.

**IMPORTANCE:** Spaceflight and spaceflight-analogue conditions alter bacterial physiology in unexpected ways that are important for pathogenesis, yet the mechanisms by which bacteria sense and respond to low fluid shear environments remain incompletely understood. This study shows that low shear modeled microgravity (LSMMG) enhances *Salmonella Typhimurium* motility and produces unexpected motility phenotypes in mutants lacking Hfq or the flagellar master regulator FlhDC. These findings indicate that flagellar biosynthesis contributes to LSMMG-enhanced motility but is not strictly required for motility under these conditions. We also suggest that flagella influence the magnitude of selected stress and infection phenotypes rather than serving as an absolute requirement for LSMMG responsiveness. Together, these results highlight the complexity of bacterial mechanotransduction under simulated microgravity conditions and advances our understanding of how a foodborne pathogen adapts to physiological low fluid shear environments encountered both in space and during terrestrial infection of the intestinal tract.

## INTRODUCTION

Microorganisms sense and respond to physical forces in their environment, including fluid shear and surface-associated forces ^1–6^. The quiescent, low fluid shear environments present in both spaceflight and LSMMG culture reflect physiological fluid forces experienced by pathogens in the infected host (*e.g.,* between the intestinal brush border), and have been shown to alter cellular processes in bacteria that shape their interactions with both the host and the environment ^1,7–13^. Indeed, culture of *S*. Typhimurium under these conditions has been associated with increased virulence in animal models, enhanced *in vitro* colonization of human tissues, increased resistance to pathogenesis-related stressors, enhanced biofilm formation, and global transcriptomic and proteomic reprogramming ^9,14–18^. Despite these well-documented phenotypic changes, the cellular mechanisms by which *Salmonella* senses and responds to spaceflight and LSMMG conditions remain incompletely defined. One prevailing hypothesis regarding how *Salmonella* and other bacteria perceive spaceflight-associated alterations in fluid dynamics posits that they do so through cell-surface structures that directly interface with the extracellular environment. Many motile bacteria, including *S*. Typhimurium, possess flagella, which are among the largest and most mechanically active appendages on the bacterial cell surface and are classically associated with motility and chemotaxis, as well as host colonization ^6,19,20^. Beyond their role in propulsion, flagella have been implicated in surface sensing, mechanotransduction, and the regulation of virulence-associated pathways in diverse bacterial species ^6,21–23^. Because flagellar assembly and operation are energetically costly, their expression and activity are tightly regulated in response to environmental and physical cues, consistent with a hypothetical model in which flagella function as key mechanosensory structures capable of transducing spaceflight- and LSMMG-associated physical signals into downstream regulatory responses ^6,13,24–26^.

The first global transcriptomic analyses of bacteria cultured under LSMMG conditions revealed altered expression of *S*. Typhimurium genes involved in flagellar biosynthesis, motility, and chemotaxis, representing the earliest evidence implicating flagella-associated pathways in LSMMG responses ^15^. Subsequent transcriptomic and proteomic profiling of *S*. Typhimurium across multiple spaceflight missions and LSMMG studies have reinforced these findings ^7,9,15–17^. Changes in motility-associated genes have also been repeatedly observed following these initial reports across multiple bacterial species, spaceflight analogue experimental platforms, and spaceflight missions, albeit with context-dependent variation in both magnitude and direction of expression ^7,9,15–17^. Moreover, Kim et al. observed that flagellar-driven motility was essential for the formation of a unique biofilm architecture by *Pseudomonas aeruginosa* during spaceflight, further implicating flagella in the regulation of surface-associated microbial behaviors under microgravity conditions ^11^.

The repeated observations of changes in motility-associated pathways and/or motility-associated phenotypes across diverse microbes suggest that flagellar regulation under LSMMG and spaceflight conditions may be integrated within broader stress-responsive regulatory networks. Under terrestrial conditions, flagellar biosynthesis and motility in *S.* Typhimurium are influenced by several global stress response regulators, including Hfq and RpoS ^27–30^. Hfq is an RNA-binding chaperone that pairs with small non-coding RNAs (sRNA) to post-transcriptionally regulate the expression of genes involved in virulence, envelope homeostasis, metabolism, biofilm formation, and motility ^31^. Deletion of *hfq* in *S*. Typhimurium has been associated with major motility defects under conventional culture conditions ^32^. RpoS is a master response regulator that enables resistance of *S*. Typhimurium to harsh environmental conditions, including those commonly encountered by the pathogen during stationary phase (e.g., nutrient deprivation, pH stress, osmotic stress), and like Hfq, is critical for *Salmonella* virulence and pathogenesis ^33,34^. Notably, Hfq regulates translation of the *rpoS* transcript through sRNA-dependent mechanisms, functionally linking these regulatory pathways ^35–39^. Our prior spaceflight and LSMMG studies have implicated Hfq and RpoS in a complex regulatory landscape in *S*. Typhimurium that extends beyond known stress response pathways, positioning these regulators as candidate integrators of motility-associated remodeling under LSMMG conditions ^9,17,40,41^.

Although altered flagellar gene expression has been repeatedly observed under LSMMG and spaceflight conditions in *S*. Typhimurium, whether flagella themselves are directly responsible for driving these phenotypic responses or instead function indirectly as modulators to fine-tune responses within a broader LSMMG-responsive network, has not been directly tested. In the present study, we sought to better define the role of flagella in *S.* Typhimurium responses to LSMMG culture, including motility, stress adaptation, and host-pathogen interactions. The findings from this study provide new insight into the mechanistic basis of bacterial adaptation to LSMMG culture and advance our understanding of microbial mechanotransduction under spaceflight-analogue conditions.

## MATERIALS AND METHODS

### Bacterial culture in the RWV

Bacterial strains (**Table 1**) were grown to stationary phase under LSMMG and 1x*g* control conditions using Rotating Wall Vessel (RWV) bioreactors, a spaceflight-analogue suspension culture system, or under conventional shaking flask conditions **(Figure 1)**. For each experiment, single colonies were isolated from frozen stocks streaked onto LB agar, and five independent colonies were inoculated into 5 mL Lennox broth (LB). Cultures were incubated overnight at 37 °C for 15 h with aeration at 250 rpm. Overnight cultures were then diluted 1:200 into fresh LB and aseptically loaded into two RWV vessels and air bubbles were carefully removed prior to incubation. RWV cultures were maintained in either the LSMMG or 1x*g* orientation at 25 rpm and grown for 24 hr at 37 °C **(Figure 1)**. For select studies, flask cultures were performed as described above with the exception that they were grown for 24 h at 37 °C. Following incubation, cultures were immediately unloaded from RWV vessels or flasks and used for downstream assays.

**Table 1.** Bacterial strains used in this study.

| Strain name | Description | Reference |
| --- | --- | --- |
| χ3339 | Wild-type *Salmonella* Typhimurium *rpsL hisG* (SL1344 animal passage) | Gulig and Curtiss<sup>42</sup> |
| $\Delta flhDC$ | TH3730. *P<sub>flhDC</sub>5451::Tn10dTc[del-25]* | Frye *et al*<sup>43</sup> |
| $\Delta rpoS$ | χ3339 $\Delta rpoS$. The *rpoS* gene (996bp) was deleted by conjugation using a suicide vector | Franco *et al*<sup>41</sup> |
| $\Delta hfq$ | χ3339 $\Delta hfq$. The *hfq* gene (309bp) was deleted by conjugation using a suicide vector | Barrila *et al*<sup>17</sup> |
| SL1344 | Wild-type SL1344 | Hoiseth *et al*<sup>44</sup> |

**Figure 1.**
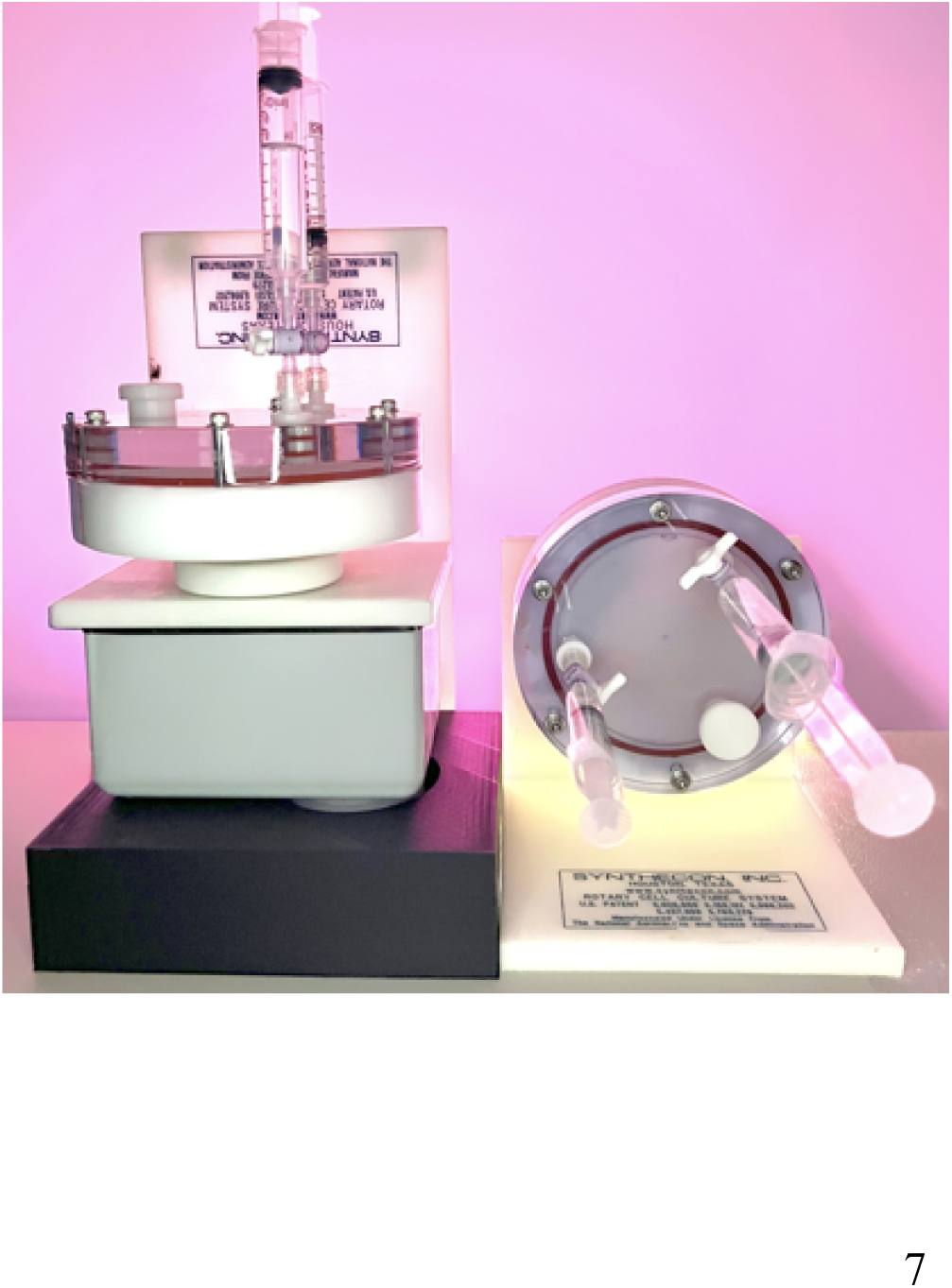
Rotating wall vessel bioreactor culture conditions. Representative image of a rotating wall vessel (RWV) bioreactor used as a spaceflight culture analogue, showing the two bioreactor orientations used in this study. Overnight cultures were diluted 1:200 in fresh LB medium, loaded into separate RWV bioreactors, and cultured at 25 rpm in either the LSMMG or 1 × *g* orientation. In the LSMMG orientation, solid-body rotation of the medium maintains bacterial cells in suspension under low-fluid-shear conditions, modeling aspects of the spaceflight environment. In the 1 × *g* orientation, bacterial cells sediment to the bottom of the vessel during rotation, thereby disrupting the low-fluid-shear condition.

### Motility assays

Overnight cultures of each strain were diluted 1:200 in fresh LB and loaded into RWVs or a flask as indicated, and grown for 24 hours at 37 °C. Following incubation, cultures of each strain were spotted onto soft agar plates (0.35% agar, 0.5% NaCl, 1% tryptone) and incubated at 37°C until the motility zones were sufficiently large to be measured. Faster motility strains, (WT and Δ*rpoS*), were incubated for 4 to 6 hours, while strains with lacking motility (Δ*hfq* and Δ*flhDC*) were incubated for 9 to 13.5 hours. Photographs of each plate were taken for analysis. Differences in motility zone areas between LSMMG and 1x*g* conditions were assessed using ImageJ. The plate diameter was set to 600 pixels/inch for consistent measurement. To decrease observational bias, experiments were performed multiple times by multiple individuals. Statistical analysis was performed using Prism 10, with paired tests applied. Area ratios (LSMMG/1x*g*) were calculated using the median values.

### Supernatant exchange and motility assay

Culture supernatants were collected by centrifugation (7,000 rpm, 7 min) to remove bacterial cells and subsequently filter-sterilized using 0.2-µm pore-size filters. Cell pellets were resuspended in the conditioned test supernatant, as indicated in the Results section, and immediately used for motility analysis, as described above. Negative controls were included to confirm the absence of viable cells, by spot-inoculating filter-sterilized, cell-free supernatants onto soft agar plates and monitoring for growth.

### Transmission Electron Microscopy (TEM)

TEM was performed to validate the presence and absence of flagella on the WT and *flhDC* mutant, respectively. Bacterial cultures were fixed without prior centrifugation by adding fixative directly to the culture. Briefly, 1 mL of bacterial culture was mixed with 4 mL of fixative solution consisting of 2% (v/v) glutaraldehyde and 2% (v/v) formaldehyde prepared in phosphate-buffered saline (PBS) or 0.2 M cacodylate buffer. Samples were fixed overnight at 4 °C. Following fixation, cells were washed three times with Dulbecco’s phosphate-buffered saline (DPBS) and allowed to gravity settle to prevent damage to flagella. Carbon-coated copper grids were plasma cleaned for approximately 1 min prior to sample loading to improve surface hydrophilicity. Fixed bacterial suspensions (5–8 µL) were applied to the smooth side of each grid and allowed to adsorb for 2 min. Excess liquid was removed by gentle blotting using filter paper. Grids were then negatively stained by placing the bacterial side of the grid in contact with staining solution, followed by blotting to remove excess stain. This staining step was repeated up to three times as needed. After staining, grids were air-dried for approximately 2 min. Prepared grids were imaged using a Talos L120C transmission electron microscope (Thermo Fisher Scientific) operated under standard imaging conditions.

### Pathogenesis-related stress assays

*S.* Typhimurium WT SL1344 and Δ*flhDC* were cultured in RWVs as described above. Following RWV growth, cultures were normalized and subjected to individual stress conditions as described previously ^2^. For acid stress, bacterial suspensions were exposed to acidified LB adjusted to pH 3.5 with 1M citrate buffer. For bile stress, cultures were incubated in LB supplemented with 10% (w/v) bile salts. For thermal stress, bacterial cultures were exposed to 52 °C. At indicated time points, samples were serially diluted in PBS and plated onto LB agar for enumeration of colony-forming units (CFU). Plates were incubated at 37 °C, and viable CFU were quantified to assess bacterial survival. Percent survival was calculated by normalizing CFU recovered following stress exposure to the initial inoculum. All stress assays were performed using three independent biological replicates. Statistical significance between conditions was assessed using unpaired two-tailed Student’s t-tests, with a significance threshold of p < 0.05.

### Infection assays

Human colonic epithelial cells (HT-29; ATCC HTB-38) were cultured as monolayers in GTSF-2 media as previously described ^41^. Cell culture media was removed about 30 mins prior to infection and 250uL of fresh media was added to each well to reduce media volume and enhance contact of bacteria-host cells. A gentamicin protection assay was then performed as described previously ^16^. Briefly, WT SL1344 and Δ*flhDC* mutant bacteria were added at a multiplicity of infection (MOI) of 1 – 10 bacteria to host cells. Samples were collected at 30 min, 3 h, and 24 h. For each time point, host cells were washed 2-3 times with HBSS, lysed with 0.1% sodium deoxycholate, and serial dilutions of the lysate plated on LB agar to assess CFU per mL. Gentamicin was added after the 30 min (50 μg/mL) and 3 h (10 μg/mL) time points in order to eliminate remaining extracellular bacteria. Percent survival was calculated by normalizing the CFU/mL at each time point to the initial bacterial inoculum. *E. coli* HB101 was used as a non-invasive control strain. Each experiment represents two or three biological replicates (as indicated in the figure legend) with three technical replicates per condition. The mean of three CFU counts were considered as one sample for statistical calculations. Data from 9 samples were analyzed with two-way ANOVA with 95% confidential interval (alpha = 0.05). Multiple t-tests were performed to compare between groups under LSMMG or 1x*g* conditions (alpha=0.05).

## RESULTS

### LSMMG culture enhances motility of wild-type S. Typhimurium and the rpoS and hfq isogenic mutants

Prior studies have reported altered expression of flagellar genes in *Salmonella* during spaceflight and LSMMG conditions ^9^. To test whether these molecular changes translate into functional changes in flagellar motility, we performed soft-agar swimming assays on wild-type *S.* Typhimurium χ3339 following culture under LSMMG and reoriented 1x*g* conditions in the RWV. Wild-type *S*. Typhimurium exhibited a slight, but significant increase in motility following LSMMG culture relative to the control (**Fig. 2A**). This finding is consistent with our previous gene expression studies using this strain, which showed upregulation of motility and chemotaxis genes under LSMMG during stationary phase ^17^.

**Figure 2.**
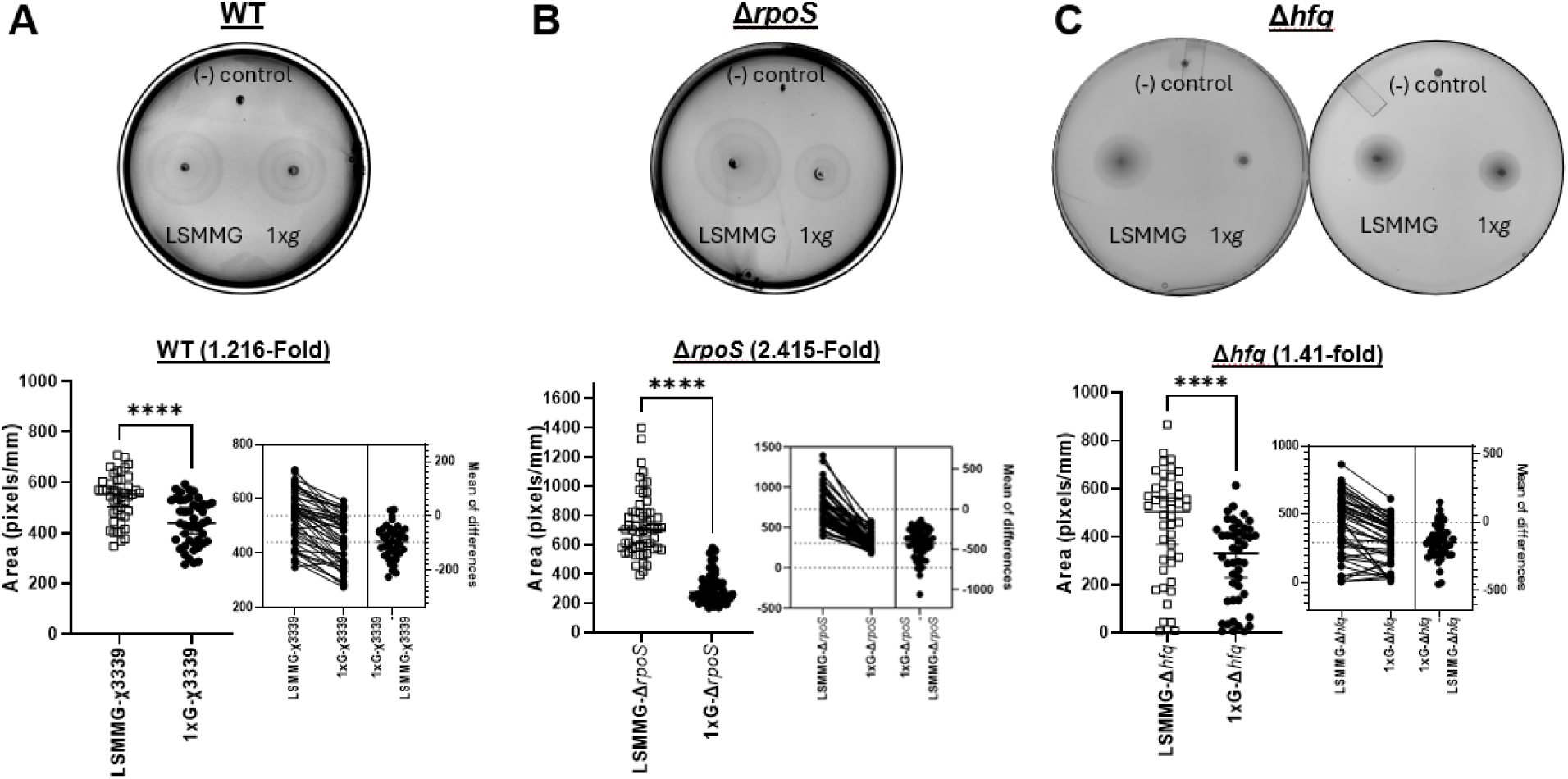
Quantitative analysis of bacterial motility on soft agar plates. Bacterial motility was tested on soft agar plates (see section methods for detail). **A – C (top).** Representative image of each strain from motility assay. **A – C (bottom).** Paired t-test results indicated a significant difference between LSMMG vs 1xg conditions for each strain with p<0.0001. Bars indicate the median values. The area ratios (LSMMG/1x*g*) are indicated in parentheses. The negative (-) control was the flask-cultured Δ*flhD/C* strain.

To identify potential regulatory determinants underlying these functional changes, we examined whether LSMMG-enhanced motility in *S*. Typhimurium required global stress response regulators RpoS and Hfq. Interestingly, similar to the WT strain, both isogenic Δ*rpoS* and Δ*hfq* mutants exhibited enhanced swimming motility following LSMMG culture (**Fig. 2B-C**). The finding that the Δ*hfq* mutant was motile following RWV culture was highly unexpected, as this mutant is typically non-motile under traditional shaking culture conditions ^32^. Paired t-test analysis revealed a significant difference between LSMMG and 1x*g* conditions for each mutant: a 2.415-fold increase (95% CI: 2.052, 2.846) in the Δ*rpoS* mutant and a 1.51-fold increase (95% CI: 1.139, 1.742) in the Δ*hfq* mutant (**Fig. 2B-C**). These findings suggest that LSMMG culture enhances the motility of *S*. Typhimurium even in the absence of RpoS and Hfq.

### LSMMG alters motility through both flagella-dependent and independent mechanisms

Given the essential role of flagella in bacterial swimming motility, we evaluated RWV-associated swimming behavior of a flagella-deficient Δ*flhDC* mutant, which lacks the master transcriptional regulator of flagellar biosynthesis ^43^. Unexpectedly, despite our prediction that this mutant would be non-motile, motility was observed following both LSMMG and 1x*g* culture, whereas the mutant remained non-motile when grown in traditional flask cultures (**Fig. 3A**). Together, these findings suggest that RWV culture conditions can enable motility in the flagella-deficient mutant strain through flagella-independent mechanisms. Furthermore, the Δ*flhDC* mutant exhibited a reversal in the relative motility trends between LSMMG and 1xg compared to the wild-type, with higher motility observed for the 1x*g* cultures (**Fig. 2 and 3**). This pattern indicates that flagella modulate the relative motility of *S*. Typhimurium to LSMMG versus 1x*g* conditions. TEM analysis validated the absence of flagella in the Δ*flhDC* mutant (**Fig. 3C**), indicating that the observed motility arises from flagella-independent mechanisms, such as adhesin-mediated surface-associated movement. Taken together, these findings suggest that while flagella influence LSMMG-associated motility, additional, uncharacterized regulatory mechanisms support motility in the RWV in the absence of flagella.

**Figure 3.**
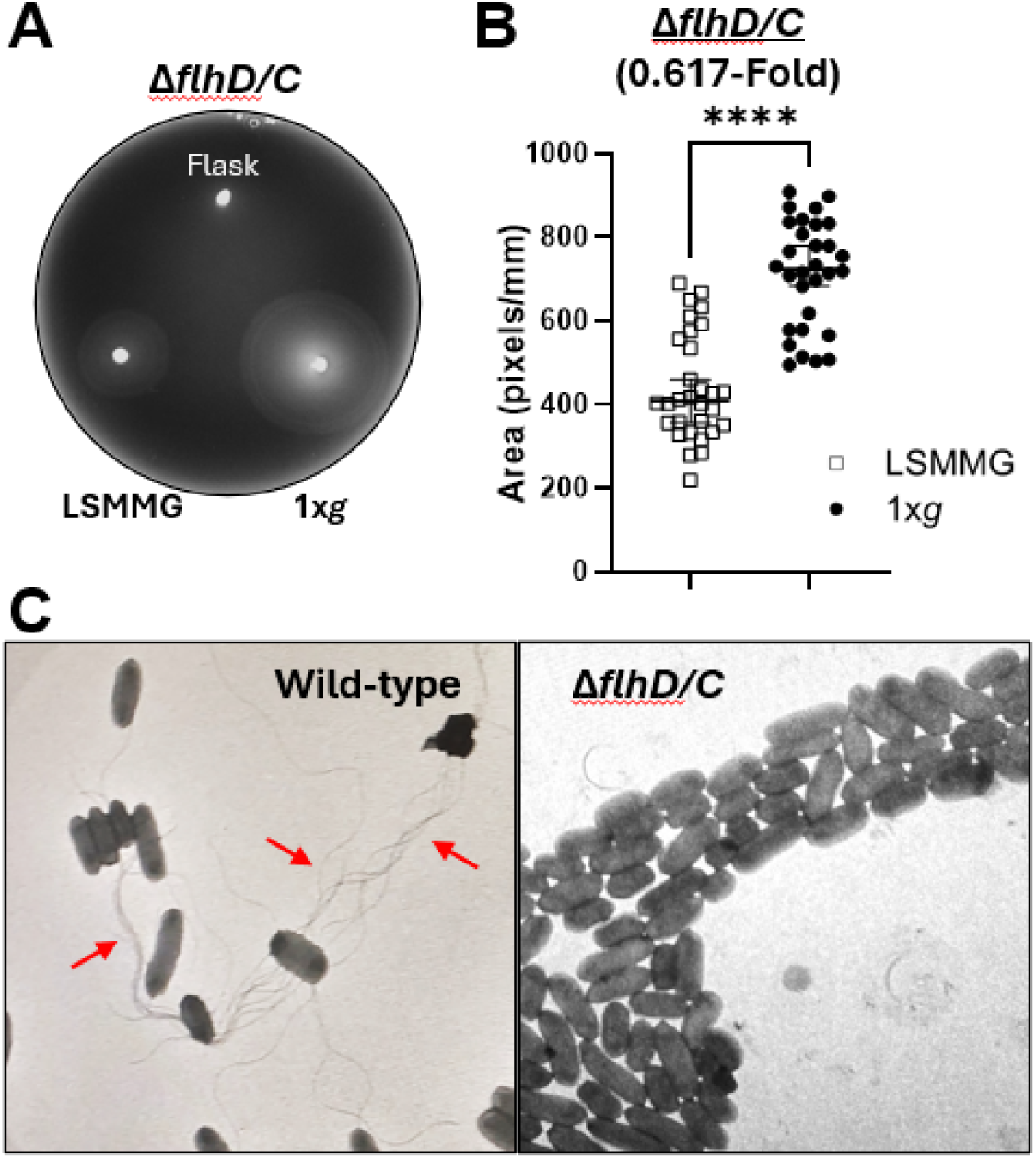
Quantitative assessment of bacterial motility and ultrastructural confirmation of flagellar loss in the *flhD/C* mutant. **A.** Photograph of the motility assay. Cultures were grown in RWVs under LSMMG or 1x*g*, or in shaking flasks and motility assessed on 0.3 % soft-agar plates. **B.** Comparison of motility between LSMMG and 1x*g* conditions. Paired t-test (n=30), p < 0.0001 (t, df=12.67, 29). Bars represent median values. The area ratios (LSMMG/1x*g*) are indicated in parentheses. **C.** Transmission electron microscopy (TEM) images showing flagella (arrows) in the wild type (left) and absence of flagella in the *flhD/C* mutant (right).

### Conditioned supernatant exchange does not alter LSMMG-associated motility phenotypes

To determine whether these flagella-independent motility phenotypes could be mediated by secreted extracellular factors, we examined whether the exchange of conditioned supernatants derived from different Δ*flhDC* cultures (i.e., LSMMG, 1x*g*, or flask) was sufficient to alter motility. Following culture under their respective conditions, bacterial cells were pelleted by centrifugation and the supernatants were filter-sterilized to remove residual cells. The cell pellets and filtered supernatants were then used for cross-mixing experiments. First, we tested whether supernatants from LSMMG or 1x*g* conditions could restore the motility of flagella mutants grown in flask conditions by mixing flask-grown bacterial cell pellets with supernatant from either LSMMG or 1x*g* conditions (**Fig. 4B**). Conversely, we also investigated whether supernatants from flask conditions inhibited motility, by mixing LSMMG- or 1x*g*-grown cell pellets with supernatants from flask conditions (**Fig. 4C**). Our results indicated that switching supernatants did not alter the motility pattern: flask-grown cells remained non-motile even in the presence of supernatant from RWV cultures, while RWV-grown cells retained their motility pattern, with enhanced motility in 1x*g* conditions compared to LSMMG. These findings suggest that the RWV-induced motility alterations are not attributable to soluble factors present in the supernatant (**Fig. 4**).

**Figure 4.**
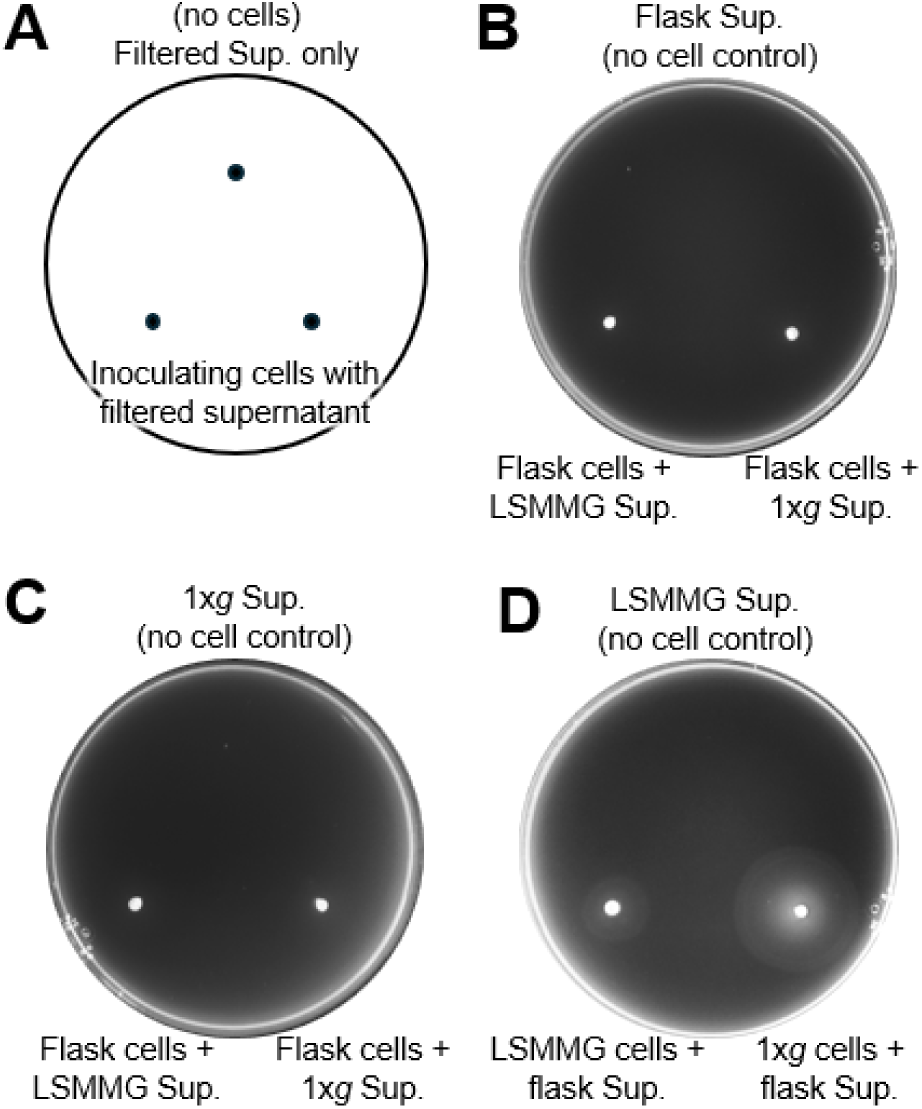
Supernatant-swapping motility assay of the flagellar mutant (Δ*flhD/C*) on soft agar. Motility of flagella mutant was assessed on 0.3% soft agar plates. **(A)** Inoculation scheme used for all plates. The top spot contains only the cell-free, filtered culture supernatant (no cells) and serves as a control to confirm the absence of viable bacteria in the supernatant preparation. The bottom two spots contained washed bacterial cells that were centrifuged and resuspended in the indicated supernatants prior to inoculation. ‘Sup’ refers to supernatant. **(B) Top spot**: cell free, filtered supernatant from flask-grown culture. **Bottom two spots**: Flask-grown Δ*flhD/C* cells resuspended in either LSMMG-derived or 1x*g*-derived supernatant. **(C) Top spot**: cell free, filtered supernatant from 1×g-grown culture. **Bottom two spots**: Flask-grown Δ*flhD/C* cells resuspended in either LSMMG-derived or 1x*g*-derived supernatant, replicating the bottom spots in panel B. **(D) Top spot**: cell-free, filtered supernatant from 1×g-grown culture. **Bottom two spots**: RWV-grown Δ*flhD/C* cells in either LSMMG or 1x*g* condition resuspended in supernatant derived from flask-grown cultures.

### Flagella modulate stress resistance and infection phenotypes following RWV culture

To assess the contribution of flagella to pathogenesis-relevant stress adaptation following RWV culture, we subjected LSMMG- and 1x*g*-cultured WT and isogenic Δ*flhDC* mutant *S.* Typhimurium to acid (pH 3.5), bile salts (10%), and thermal stress (52 °C). Both WT and *ΔflhDC* mutant 1x*g* cultures exhibited higher resistance to acid and bile stressors compared to LSMMG cultures (**Fig. 5A-B**). Trends for the wild-type were consistent with previous studies ^9,14,41^. While the loss of flagella did not alter the direction of the stress responses, a comparison of the survival ratios (1x*g*/LSMMG) revealed a consistently reduced magnitude between conditions for the mutant relative to WT (**Fig. 5A-B**). This is consistent with a limited contribution of flagella to these LSMMG-associated stress responses in *Salmonella*. In contrast, thermal stress response profiles revealed a more pronounced effect. While there was no significant difference in resistance between LSMMG and 1x*g* cultures of the WT strain, LSMMG cultures of the *ΔflhDC* mutant exhibited significantly enhanced survival relative to 1x*g* controls (**Fig. 5C**). Thus, unlike acid and bile stress, thermal stress revealed an LSMMG-associated survival advantage specifically in the flagella-deficient background, indicating a stressor-specific role for flagella in modulating LSMMG-associated thermal stress adaptation.

**Figure 5.**
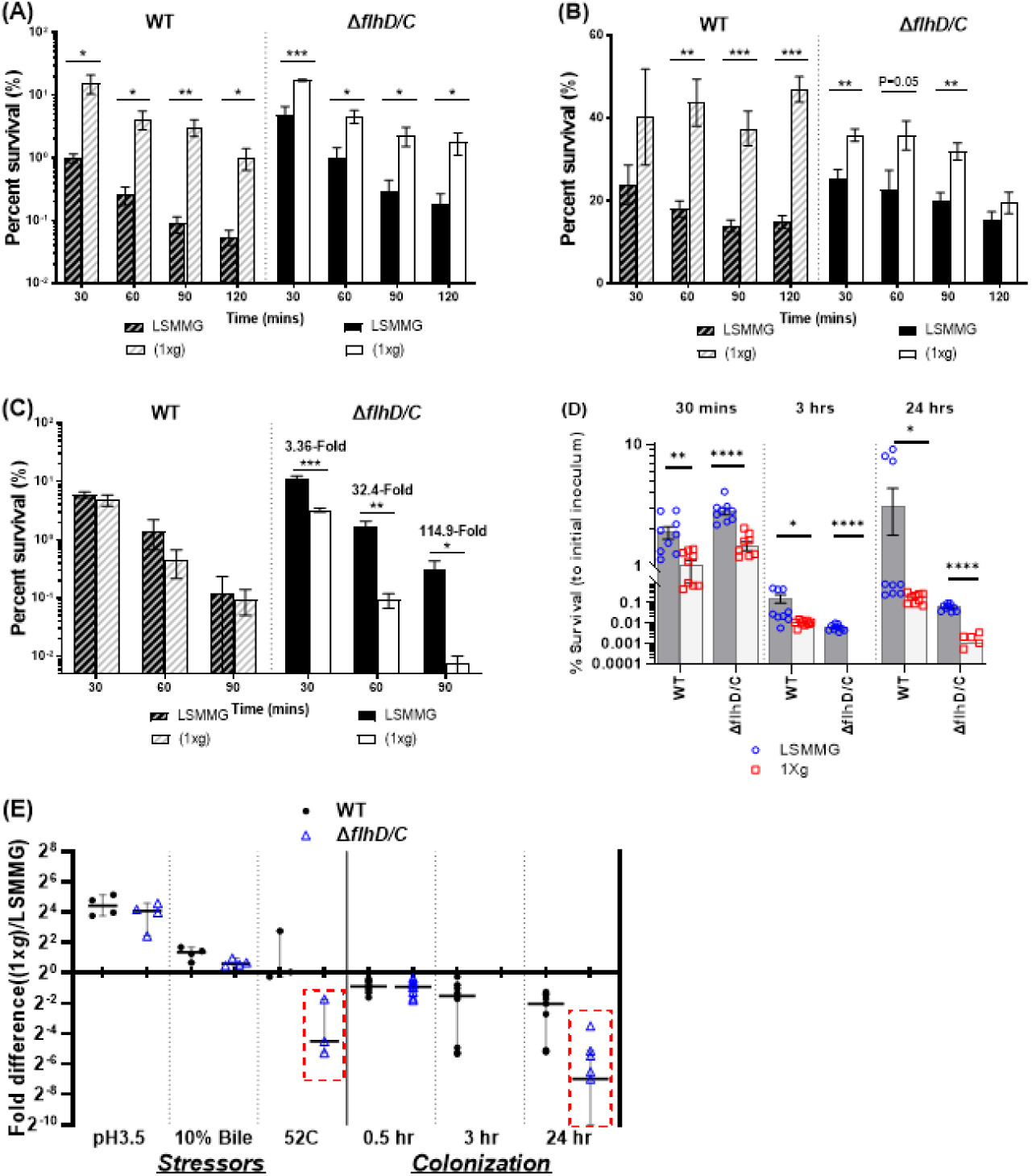
Stress response and host colonization of *S.* Typhimurium flagella mutant compared to the wild type. The wild-type (WT) *S. Typhimurium* and its isogenic flagella-deficient mutant (Δ*flhD/C*) are tested.**(A–C)** Stress response assays under different pathogenesis-related conditions: **(A)** acid stress (pH 3.5), **(B)** 10% bile salt exposure, and **(C)** thermal stress (52 °C). Data represent mean ± SEM from three independent biological replicates, each with two to three technical replicates. Statistical significance was determined using two-tailed *t*-tests (**p < 0.01; \**p < 0.001).* **(D) *Host colonization assay****. S. Typhimurium wild-type and the ΔflhD/C* mutant were cultured under low-shear modeled microgravity (LSMMG) or 1×*g* conditions and used to infect HT-29 monolayers. Samples were collected at 0.5 h, 3 h, and 24 h post-infection, and viable bacterial counts (CFU/mL) were determined. Percent survival was calculated relative to the initial inoculum. Data represent mean ± SEM (n = 9). Two-way ANOVA (α = 0.05) followed by multiple *t*-tests was performed to compare groups (*p < 0.05; **p < 0.01; ***p < 0.001; ****p < 0.0001). **(E)** Fold difference between fluid shear conditions. The ratio of survival under 1×*g* to LSMMG conditions was calculated for each group. Median values and 95% confidence intervals are shown. The red box highlights regions of interest described in the main text.

To assess whether flagella play a role in the infection of human intestinal epithelium, we performed a gentamicin protection assay to assess the impact of the Δ*flhDC* mutation on adherence (0.5h), invasion (3h) and intracellular survival (24h) relative to the wild-type in HT-29 monolayers. Consistent with our previous findings (Barrila 2022; Franco), WT LSMMG cultures exhibited enhanced adherence, invasion, and intracellular survival/replication relative to 1x*g* cultures. While a similar pattern of LSMMG-enhanced colonization was also observed for the *ΔflhDC* mutant (**Fig. 5D**), the wild-type bacteria consistently exhibited higher colonization at all infection stages (0.5 h, p = 0.0011; 3h p = 0.0351; 24h p = 0.0250). Moreover, two-way ANOVA revealed a significant interaction between the RWV culture environment and flagellar status at 24 h post-infection (p = 0.0407), indicating that the presence of flagella in the WT amplifies the magnitude of LSMMG-dependent intracellular survival rather than determining the direction of the response. These data demonstrate that LSMMG culture and flagella act in concert to influence aspects of *Salmonella* pathogenesis, particularly during the intracellular phase of infection. Fold-difference analysis (1x*g*/LSMMG) revealed that LSMMG-dependent responses are stressor-specific in direction, favoring 1x*g* for acid and bile stress but LSMMG for thermal stress and infection (**Fig. 5E**). Collectively, these findings demonstrate that loss of flagella selectively alters the magnitude of these responses, with the strongest effects observed during thermal stress and intracellular survival (**Fig. 5**).

## DISCUSSION

In this study, we assessed the impact of LSMMG culture on the swimming motility of *S*. Typhimurium and evaluated whether flagella serve as key mechanosensory structures in the motility, stress, and infection responses of the pathogen to this environment. We also investigated potential roles of the global stress response regulators RpoS and Hfq in regulating motility in these environments. Under the conditions of this study, LSMMG culture enhanced the motility of wild-type *S*. Typhimurium, with this heightened motility retained in Δ*rpoS* and Δ*hfq* backgrounds. These findings were consistent with our prior transcriptomic analyses performed in the RWV under identical culture conditions (i.e., media, temperature, growth phase) which demonstrated increased expression of numerous motility genes for both the wild-type and Δ*hfq* mutant under LSMMG relative to 1x*g* ^17^. Nevertheless, phenotypic validation that the Δ*hfq* mutant was indeed motile following both LSMMG and 1x*g* culture and exhibited enhanced motility under LSMMG was particularly notable, as this mutant is typically non-motile under conventional shaking culture conditions^32^ (**Fig. 3A**).

One potential explanation for the enhanced motility of the *hfq* mutant following LSMMG culture relates to the quiescent, low fluid shear conditions present within the RWV. Mathematical modeling and/or dye tracer mixing studies have shown that LSMMG culture allows cells to remain suspended in a physiological low fluid shear environment relevant to that encountered by *Salmonella* between the intestinal brush border epithelium, while conversely, 1x*g* cultures display more rapid mixing with eventual settling of the cells at the bottom of the reactor, where they are expected to encounter frictional forces along the membrane of the RWV prior to full sedimentation ^16,45,46^. Deletion of *hfq* in *Salmonella* is associated with disruption of sRNA-mediated control of outer membrane biogenesis and stress responses, leading to a range of pleiotropic effects, including chronic envelope stress associated with dysregulated porin expression, impaired periplasmic proteostasis, and reduced membrane integrity ^32^. Accordingly, these envelope-associated defects would be expected to increase mechanical sensitivity of the mutant, particularly under conditions of elevated fluid shear and surface-associated friction, such as those encountered in conventional shaking cultures and, to a lesser extent, in 1x*g* RWV cultures. Under LSMMG, the reduction in fluid shear stress may partially alleviate envelope-associated mechanical strain, thereby permitting observable motility phenotypes that are otherwise suppressed in high shear conditions, even in the absence of *hfq*.

Unexpectedly, the *ΔflhDC* mutant also exhibited motility following RWV culture, despite lacking flagella. This motility pattern differed from that of the wild-type, with greater motility under 1x*g* than LSMMG conditions. Supernatant exchange experiments with this mutant confirmed that these trends were due to cell-intrinsic properties rather than secreted extracellular factors. Collectively, these observations indicate that RWV-associated motility can arise in the absence of flagellar biosynthesis and therefore reflects engagement of flagella-independent motility mechanisms. The persistence of motility in the mutant suggests the involvement of alternative surface-associated motility processes, potentially including gliding, twitching motility mediated by type IV pili, or other non-flagellar movement mechanisms, which warrants further investigation. Thus, within the RWV environment, flagella are not an absolute requirement for motility but instead act as modulators that shape the magnitude and directional pattern of the response.

Pathogenesis-related stress and infection assays comparing wild-type and the Δ*flhDC* mutant revealed that the overall direction of LSMMG-dependent phenotypic shifts was preserved between the two strains for most phenotypes, indicating that these responses do not strictly require flagellar biosynthesis. In contrast, thermal responses exhibited divergent LSMMG dependence between the wild-type and mutant, with the LSMMG-cultured mutant exhibiting enhanced thermal stress resistance, coinciding with conditions in which motility behavior also diverged. Given flagellar motility and flagellar regulatory networks are known to respond to temperature and other host-associated environmental cues, this divergence raises the possibility of an interaction between LSMMG-responsive thermal adaptation pathways and flagella-associated regulatory processes ^47,48^. Similarly, although LSMMG enhanced host cell adherence, invasion, and intracellular survival in both wild-type and flagella-deficient strains, the magnitude of this enhancement, particularly during late intracellular survival, was influenced by the presence or absence of flagella. Although the mechanisms underlying this modulation remain to be defined, prior evidence linking flagellar regulation to SPI-1 regulatory circuits provides one possible framework for interpreting how flagella may tune the magnitude of LSMMG-enhanced infection phenotypes ^49–51^. Collectively, these findings indicate that flagella regulate the magnitude, but not the overall directional trend, of most LSMMG-dependent stress and infection phenotypes, supporting a role for flagella as mechanical modifiers that tune the intensity of LSMMG responses rather than serve as primary sensors of these outcomes.

Taken together, our results support a model in which *S.* Typhimurium integrates cues associated with LSMMG culture through multiple regulatory pathways that extend beyond canonical stress regulators and flagellar biosynthesis. In this framework, rather than serving as obligate mechanosensors, flagella may function as modulatory components within a broader LSMMG-responsive network that coordinates motility, stress adaptation, and infection-relevant phenotypes. Importantly, the persistence of motility and LSMMG-dependent phenotypes in the absence of flagella underscores the contribution of additional, flagella-independent mechanisms to these responses. These findings expand our understanding of bacterial adaptation to physiological low fluid shear environments and highlights the complexity of bacterial mechanosensing, suggesting new avenues for exploring how LSMMG-associated environmental changes shape microbial physiology and pathogenesis in extreme environments.

## Author Contributions

J.Y. and J.B. contributed to conceptualization, supervision, data analysis and interpretation, and writing of the original draft. J.Y., K.P.F.M., L.B., C.C., B.Y.K., R.R.D., and S.G. performed experiments and contributed to data analysis. C.M.O. contributed to data interpretation, discussion, and manuscript review and editing. RJCM. provided materials and contributed to manuscript review and editing. C.A.N. contributed to investigation, project administration, data interpretation, discussion, and manuscript review and editing.

## ACKNOWLEDGEMENTS

This work was supported by NASA grants, NASA NNX09AH40G (CAN, CMO); NASA 80NSSC20K0016 (CMO, CAN, JB); NASA 80NSSC24K0744 (CAN, JB, JY, CMO); NASA NNX15AL06G (CAN, JB, JY, CMO); BAC 80NSSC22K1361 (RJC, CAN, JB, JY, CMO).

